# Edge-First Ground Reaction Force Estimation with Consumer Smartwatches

**DOI:** 10.64898/2026.07.18.739307

**Authors:** Parvin Ghaffarzadeh, Debarati Chakraborty, Koorosh Aslansefat, Ali Dostan, Yiannis Papadopoulos

## Abstract

Ground reaction force (GRF) measurement remains largely confined to instrumented laboratories, limiting longitudinal monitoring in daily life. This article presents an edge-first wearable system for estimating vertical GRF from consumer smartwatches. Two Apple Watch Series 6 devices worn at the wrist and waist stream 12-channel inertial data at 100 Hz to an iPhone, where preprocessing, storage, and inference occur locally without cloud dependence. The proposed *GRFNet-MultiScale* model is a compact temporal convolutional network with four dilated residual blocks and a global context branch. Under leave-one-subject-out evaluation on 539 stance windows from 10 healthy participants, the dual-sensor system achieved a mean Pearson correlation of 0.798 with an RMSE of 257 N, while a wrist-only configuration retained 82.5% of dual-sensor correlation. Temporal attribution remained stable across validation folds and identified early-stance wrist acceleration as the dominant reproducible signal. The system is strongest for cyclic locomotion.

## 1 Introduction

GROUND reaction force is one of the most informative signals in movement analysis. It captures how the body loads and unloads during stance and underpins applications in gait assessment, rehabilitation, return-to-sport monitoring, and cumulative load estimation [1]–[4]. In practice, however, accurate GRF measurement still depends on force plates embedded in laboratory floors. These systems remain precise but fixed, expensive, and difficult to scale beyond supervised sessions. For many real-world use cases, the missing capability is not another isolated laboratory snapshot. It is repeated measurement in daily settings, where loading changes over time rather than during a single visit.

Wearable sensing offers a practical route beyond that limitation. Commodity devices already contain inertial sensors, wireless connectivity, and enough local compute to support on-device preprocessing and inference. Prior work has shown that GRF or related kinetic signals can be estimated from inertial data with promising accuracy, especially during walking and running [5]– [10]. Yet from a wearable-computing perspective, much of that literature still leaves key translation questions underdeveloped: how data move from sensor to edge node, how privacy is handled, how subject-independent generalization is tested, and how time-series explanations are made reproducible across training splits [5]–[13].

This article therefore positions wearable GRF estimation as an edge-systems problem rather than only a biomechanics prediction problem. The system uses two Apple Watch Series 6 devices as body-worn sensor nodes and an iPhone as the edge node. The phone performs local buffering, preprocessing, storage, and inference without requiring cloud connectivity for the inference path. Figure 1 in Section 3 summarizes that full stack. The predictive core is GRFNet-MultiScale, a compact temporal convolutional model that captures both short impact-related events and longer stance-scale structure. To improve trustworthiness, the article also uses time-resolved explainability methods that identify not only which channels matter, but when they matter during stance [11]–[13].

**Fig. 1.**
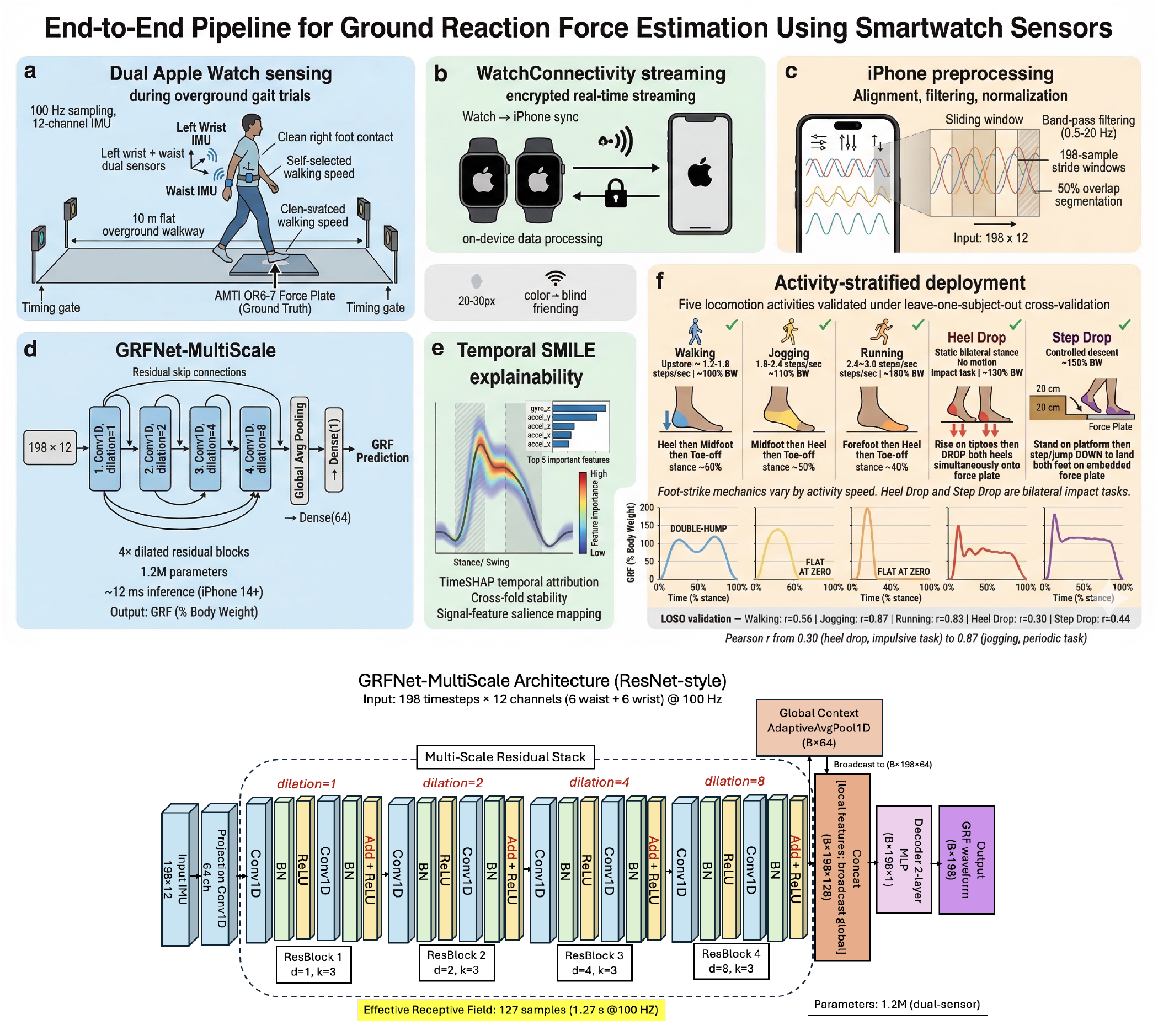
**Top:** End-to-end edge-first wearable pipeline. Two Apple Watch Series 6 devices stream 12-channel IMU data to an iPhone, which performs local low-pass filtering, alignment, normalization, GRF inference, and time-resolved explainability without cloud dependence. Activity-scoped deployment boundaries are summarised. Dataset context follows [15]; temporal explanation components are grounded in [12], [13]. **Bottom:** GRFNet-MultiScale architecture. Multi-scale dilated residual ∈*{}* blocks (*d* 1, 2, 4, 8, kernel *k*=3) process 12-channel dual-sensor IMU input. Each block uses two Conv1D layers with batch normalization, ReLU, dropout, residual addition, and post-addition ReLU. Global context from adaptive average pooling is broadcast and concatenated with local features before per-timestep decoding. Effective receptive field: 127 samples (1.27 s); parameters: 1.2M (dual-sensor) [14].

This article makes four contributions. First, it presents an end-to-end wearable edge pipeline using consumer devices and local phone-side execution. Second, it introduces a compact multi-scale temporal model that balances predictive accuracy with deployment practicality. Third, it evaluates the system under strict leave-one-subject-out validation, the appropriate setting for generalization to unseen users. Fourth, it shows that temporal attribution patterns remain stable across validation folds. The goal is not to claim that consumer wearables can replace laboratory force plates in all settings. It is to define what this class of edge-first wearable GRF system can already support, where its current limits lie, and how it should be evaluated for responsible deployment.

## 2 Related Work

Wearable estimation of ground reaction force (GRF) has attracted sustained interest because it offers a path beyond instrumented laboratory floors and toward repeated biomechanical monitoring in everyday settings [5]– 10]. Most prior work uses inertial measurement unit (IMU) data, sometimes combined with deep learning, to reconstruct force waveforms or related kinetic variables during walking or running. Dorschky et al. used convolutional neural networks (CNNs) to estimate sagittal-plane biomechanics from measured and simulated inertial data [5]. Johnson et al. predicted multidimensional GRF and joint moments from wearable accelerometer signals [6]. Pogson et al. focused on task- and step-specific GRF magnitude estimation during running and highlighted the difficulty of generalising to unseen individuals [7]. Wouda et al. and Alcantara et al. reported promising results for running-related vertical GRF estimation from IMU data [8], [9]. Mundt et al. estimated kinetics from motion-capture-derived inputs rather than from a consumer wearable sensing pipeline [10].

Taken together, these studies show that wearable signals do contain meaningful information about external loading. However, they also reveal a consistent translation gap between proof-of-concept prediction and deployable wearable computing systems. Many studies rely on a single sensor location, which limits placement flexibility in real-world use. Evidence for subject-independent generalization under strict leave-one-subject-out (LOSO) validation is uneven. LOSO is important here because it tests whether a model trained on one group of users can generalise to a new user it has never seen before. In addition, system-level details are often limited. Buffering, synchronisation, local storage, communication behavior, and end-device constraints are not always described in enough detail to assess edge deployment. Practical issues such as sensor placement sensitivity, sampling rate, and device-orientation error also remain important for wearable performance outside controlled laboratory conditions.

Table 1 positions the present article against representative studies on those translation dimensions rather than on accuracy alone. The aim is not to claim that prior work lacks predictive value. The point is that a wearable computing contribution must be judged across the full stack: sensing, communication, execution context, robustness, and interpretability.

**TABLE 1.**
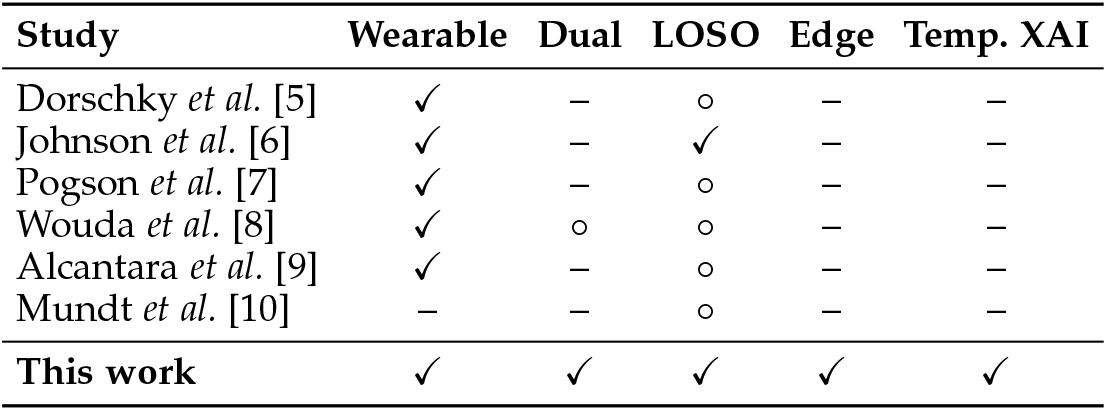
Representative wearable GRF estimation studies and key translation requirements [5]–[10]. “Edge pipeline” indicates explicit treatment of wearable-to-device communication, local processing, storage, and deployment constraints. “Temporal XAI” indicates time-resolved explanation rather than channel-only feature importance. ✓: explicitly addressed; °: partially; –: not reported.

| Study | Wearable | Dual | LOSO | Edge | Temp. XAI |
| --- | --- | --- | --- | --- | --- |
| Dorschky *et al.* [5] | ✓ | – | ◦ | – | – |
| Johnson *et al.* [6] | ✓ | – | ✓ | – | – |
| Pogson *et al.* [7] | ✓ | – | ◦ | – | – |
| Wouda *et al.* [8] | ✓ | ◦ | ◦ | – | – |
| Alcantara *et al.* [9] | ✓ | – | ◦ | – | – |
| Mundt *et al.* [10] | – | – | ◦ | – | – |
| **This work** | ✓ | ✓ | ✓ | ✓ | ✓ |

Interpretability deserves particular emphasis. In high-stakes applications, clinicians and engineers need more than a correlation score. They need evidence that the model is relying on physically meaningful signals rather than on spurious correlations [11]. This is especially important for time-series data such as gait IMU signals, where the timing of an event matters as much as the channel in which it appears. Standard feature-importance tools often compress the result into a single ranked list of channels or variables, which answers “what matters overall?” but not “what matters at which point in the step?” [12].

To address that limitation, this work uses two complementary temporal explainability methods. The first is Temporal SMILE [12], a masking-based method that perturbs short contiguous windows of a single input channel and measures how much the prediction changes. In effect, it asks: if this short segment of this signal were hidden, how much would the model’s output change? Repeating that operation across time and across channels produces a two-dimensional attribution map that shows both which signal matters and when during stance it matters. The second method is a TimeSHAP-style analysis [13], which extends SHAP-like reasoning to sequential data by assigning attribution to temporal segments rather than only to static features. This allows the stance window to be partitioned into interpretable phases and provides a complementary view of where predictive information is concentrated over time.

Using both methods is important for this article’s argument. Temporal SMILE provides an intuitive perturbation-based view of local time-channel importance, while TimeSHAP provides a segment-level attribution framework grounded in Shapley-value reasoning [13]. When both methods identify the same dominant signal pattern, the explanation becomes more credible. In this study, that agreement is then evaluated across validation folds, so the contribution is not only temporal explainability but also *reproducible* temporal explainability. That is an important distinction for deployment-oriented wearable AI, because a visually plausible explanation from a single split is much weaker than an explanation that remains stable across repeated held-out-user evaluations.

## 3 Edge-First Wearable System Architecture

The system is organized as a two-tier wearable architecture. Two Apple Watch Series 6 devices serve as body-worn sensor nodes, with one watch placed at the dominant wrist and the other at the waist. Each watch records triaxial acceleration and triaxial angular velocity at 100 Hz, yielding 12 inertial channels across the pair. The iPhone acts as the edge node. It receives the sensor streams, reconstructs them in time order, stores them locally under pseudonymized identifiers, applies preprocessing, and executes model inference. Figure 1 summarizes the end-to-end flow from on-body sensing to local prediction and activity-scoped deployment.

The end-to-end flow is as follows. First, the two watches acquire synchronized inertial measurements during locomotor activity. Second, the recorded samples are buffered on-device and transmitted to the paired iPhone through WatchConnectivity. Third, the iPhone reorders packets by sequence index, reconstructs the continuous multichannel stream, and stores the incoming data locally. Fourth, the phone applies the preprocessing pipeline, including alignment, low-pass filtering, normalization, and segmentation into fixed-length stance windows. Fifth, each processed window is passed to GRFNet-MultiScale, which estimates the vertical ground reaction force waveform for that stance phase. Finally, the predicted waveform can be used directly or reduced to derived biomechanical metrics such as impulse, peak force, and effective contact duration. In the current study, this full inference path is executed locally on the phone and does not require cloud connectivity.

This design choice is deliberate. Sending raw biomechanical data to a cloud service would increase dependency on network quality, add variable latency, and move personally identifiable motion patterns off the device. In the present architecture, the inference path does not require cloud connectivity. That improves privacy and makes the system less sensitive to connectivity variation during outdoor use. A cloud backend remains optional for model export or retraining, but it is not part of the sensing-to-inference loop evaluated here.

Communication between the watches and the phone uses WatchConnectivity. Samples are accumulated into transmission chunks of 500 samples, equivalent to 5 s of data at 100 Hz, before transfer. This choice improves robustness to transient Bluetooth drops and motion-induced attenuation because the iPhone can reorder packets by sequence index and reconstruct the stream deterministically. The tradeoff is latency. Under the current design, the system supports near-real-time monitoring rather than per-step live feedback. That tradeoff is part of the wearable-systems contribution, not an implementation footnote.

## 4 Data Collection and Preprocessing

The dataset comprises 10 healthy adults who simultaneously wore the wrist and waist watches while performing five randomised activities: walking, jogging, running, heel drops, and 20 cm step-downs. Ground-truth vertical GRF was recorded at 1000 Hz using an AMTI OR6-7 force plate and later downsampled to 100 Hz to match the inertial signals. Of 598 recorded trials, 539 passed quality screening and were retained for evaluation [15]. This cohort is modest, but the protocol is rigorous in the sense that all headline results are reported under LOSO validation rather than subject-specific tuning. The retained dataset also spans biomechanically distinct tasks, from regular cyclic gait to impulsive non-cyclic impacts, which helps define both the strengths and the limits of the system.

The preprocessing pipeline has four main stages. First, the IMU and force-plate signals are aligned using cross-correlation within a bounded temporal window. This step compensates for small acquisition offsets between the wearable devices and the laboratory force plate and ensures that each stance segment is paired with the correct force trace. Second, the IMU signals are low-pass filtered at 10 Hz and the force-plate signals at 20 Hz. These cutoffs reduce high-frequency noise while preserving the dominant structure of walking, jogging, and running. Third, stance phases are mapped to a fixed representation of 198 samples. Shorter windows are zero-padded and longer windows are truncated. Fourth, each trial is z-score normalized to reduce amplitude drift across participants and trials.

A brief standing baseline is also used to align device coordinate frames through gravity-based calibration. This matters in practice because smartwatch orientation can vary with strap placement and body segment alignment, particularly at the wrist. The calibration step does not eliminate all placement sensitivity, but it reduces gross between-trial inconsistency before the data are passed to the *×* model. The resulting input tensor has dimensions 198×12, corresponding to a fixed 1.98 s window with 12 inertial channels from the wrist-and-waist pair. This standardized representation simplifies batching during training and bounded execution during inference.

These preprocessing choices serve both modelling and deployment objectives. The fixed-size window gives the network bounded-size inputs, which simplifies memory use and execution time on the phone. The filter choices also help explain the later performance profile. The 10 Hz IMU cutoff is appropriate for robust cyclic locomotion analysis, but it attenuates some higher-frequency impact content. That is one reason the system performs much better on walking, jogging, and running than on abrupt heel-drop events. Likewise, using force-plate timing during evaluation provides precise stance segmentation for benchmarking, but it also highlights a remaining deployment gap: a fully wearable-only system will need reliable IMU-based stance detection.

From a systems perspective, the preprocessing chain is intentionally conservative. It uses operations that are lightweight, deterministic, and straightforward to port to a mobile execution environment. That matters because the deployment question is not only whether the model can predict vGRF, but also whether the surrounding signal-processing path can run consistently on an edge device without introducing hidden latency or unstable behavior. In that sense, the preprocessing pipeline is part of the contribution rather than a routine implementation detail [15].

## 5 Model Design

GRFNet-MultiScale is a compact one-dimensional temporal convolutional network designed for vertical ground reaction force estimation under edge-execution constraints. The model takes a fixed-length 198 *×* 12 input tensor derived from the wrist-and-waist inertial streams and maps it to a per-timestep vGRF prediction. The architecture is built around four dilated residual blocks with dilation rates *d*∈ *{*1, 2, 4, 8} and kernel size 3, followed by a global context branch and a lightweight per-timestep decoder. In the dual-sensor configuration, the model contains approximately 1.2 million trainable parameters. A wrist-only version reduces this to about 605 thousand parameters. Figure 1 shows the full architecture.

The architectural choice is motivated by the temporal structure of stance-phase biomechanics. A single stance window contains events at markedly different scales: heel-contact transients occur over a short interval, while loading response, mid-stance progression, and push-off evolve over longer portions of the step. A useful model must therefore represent both local and broader temporal dependencies without becoming too large for practical deployment. GRFNet-MultiScale addresses this by stacking dilated convolutions across four scales, allowing the receptive field to expand efficiently while preserving a compact parameter budget. Each residual block contains two

Conv1D layers with batch normalization, ReLU activation, and dropout (0.15), followed by residual addition. This design supports stable optimization while retaining low-level temporal information as the network builds progressively wider temporal context. The resulting effective receptive field is 127 samples (1.27 s), which is sufficient to cover the loading and mid-stance portions of all five locomotor tasks studied here [14].

The model also includes a global context mechanism to complement the local convolutional pathway. Adaptive average pooling compresses the full 198-sample window into a summary representation, which is then broadcast and concatenated with local temporal features before the final multilayer perceptron decoder. This addition allows the network to interpret local signal events relative to the broader shape of the stance window, rather than relying only on short-range filters. In effect, the architecture combines local event sensitivity with window-level context while remaining small enough for deterministic and bounded execution on the edge device. In the current implementation, the dual-sensor model runs in approximately 12 ms per window under GPU profiling, although phone-side execution has not yet been empirically validated.

A lightweight wrist-motion correction step is applied before inference. This is important because wrist-worn accelerometers capture not only gait-related dynamics but also arm swing, postural sway, and incidental upper-body movement that may be weakly related to foot-ground interaction. The correction computes the mean wrist acceleration over a short sliding window of approximately 0.5 s and subtracts a low-pass filtered 2 Hz component from the raw trace. The aim is to attenuate slower, unrelated motion while preserving the higher-frequency content more likely to reflect impact and ground-contact mechanics. The operation is intentionally simple, avoids additional learned preprocessing, and is consistent with real-time deployment requirements. In the original experiments, it was sufficient to stabilize wrist signals across participants without introducing noticeable phase distortion.

The architecture was also compared against a standard Transformer encoder. In this dataset, the Transformer achieved lower predictive performance and higher computational cost, with *r*=0.581 *±* 0.144, 2.1 million parameters, and about 42 ms inference time. This gap is plausible given the limited cohort size, since Transformer models typically benefit from substantially larger datasets to realize their capacity for long-range dependencies. By contrast, dilated temporal convolutions provide a stronger local-first inductive bias for structured gait signals. The ablation results further support this interpretation: removing the multi-scale dilation structure and setting all dilation factors to 1 reduced correlation from 0.798 to 0.705 and increased RMSE by 41 N, which was the largest degradation among the tested architectural changes. Taken together, these results indicate that the multi-scale convolutional design is not only computationally economical but also central to the model’s predictive performance.

## 6 Results and Discussion

### 6.1 Predictive Performance and Deployment Scope

All principal results were obtained under 10-fold LOSO validation. In each fold, the model was trained on nine participants and tested on the held-out participant. This is the appropriate evaluation setting for a wearable system intended to generalise to unseen users, because it mirrors the real deployment scenario more closely than within-subject adaptation. Under that protocol, the dual-sensor model achieved a mean Pearson correlation of 0.798 with an RMSE of 257 N. Figure 2(a) summarizes the participant-level variation. The spread across held-out users is real but bounded, which indicates that performance is not dominated by a small subset of subjects.

**Fig. 2.**
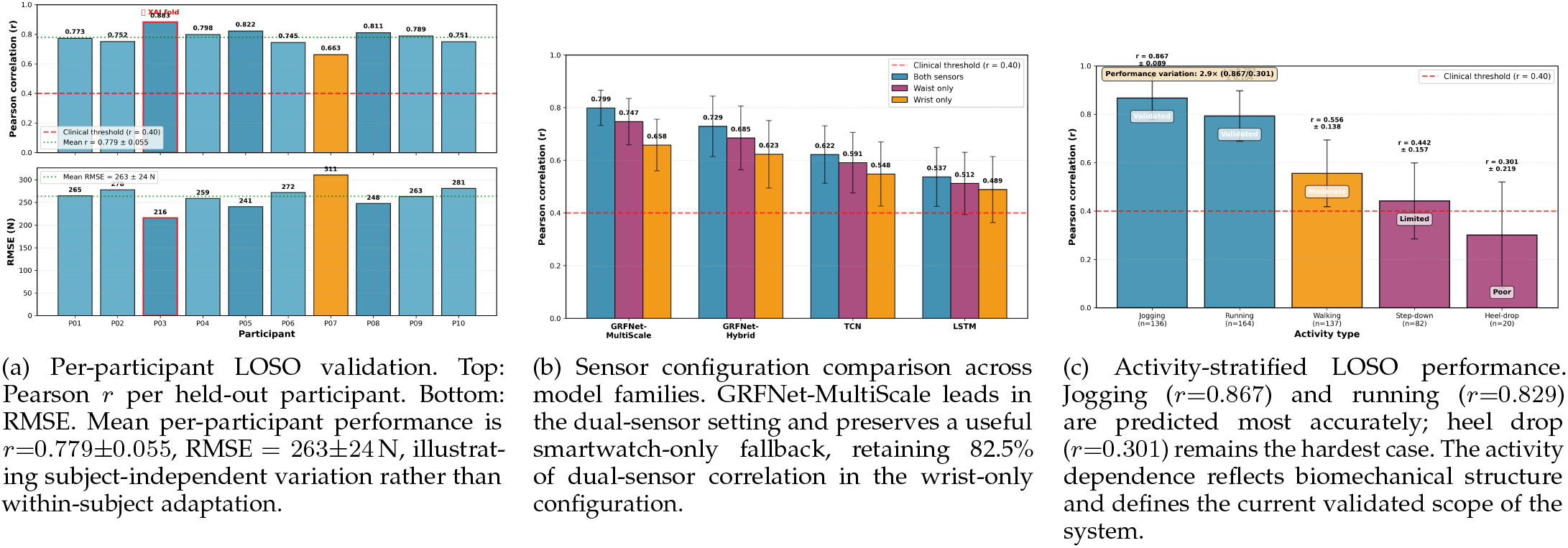
Predictive performance across evaluation dimensions under 10-fold LOSO cross-validation.

Sensor-placement flexibility matters in wearable computing, so the study also evaluates wrist-only and waist-only variants. The wrist-only configuration achieved a correlation of 0.658, retaining 82.5% of dual-sensor performance. This materially strengthens the deployment story because a standard smartwatch-only configuration remains informative when a waist device is not available. Figure 2(b) shows that this advantage persists across model families, with GRFNet-MultiScale leading the comparison.

Performance also varies by activity. Jogging and running are the strongest cases, followed by walking. Step-down performance is moderate, and heel-drop performance is low.

Figure 2(c) summarizes that activity dependence. The pattern is mechanically plausible: cyclic locomotion yields smoother and more regular GRF waveforms, whereas heel drops are brief, impulsive, and dominated by higher-frequency content that is harder to reconstruct from low-pass-filtered 100 Hz inertial signals. In practical terms, the validated deployment region is therefore cyclic locomotion, with more caution required for step-down and no present support for impulsive heel-drop-like tasks.

### 6.2 Computation and Edge Deployment Feasibility

This work reports a mean inference time of approximately 12 ms per 198-sample window on an NVIDIA A40 GPU, averaged over 50 forward passes at batch size 1. This is a useful reference measure of computational cost, but it is not yet an on-device latency measurement. The distinction matters. In a wearable system, practical latency depends not only on forward-pass time, but also on transmission, buffering, preprocessing, and phone-side execution behavior.

From an engineering perspective, the architecture is compatible with Core ML deployment in principle. The model is compact, the operations are standard, and the forward-pass cost is low enough to make phone-side execution plausible. At the same time, the present evidence does not yet support claims of fully characterized end-to-end deployment. On-device timing, energy profiling, and the effect of sustained continuous monitoring on battery life remain open. Framing that boundary clearly strengthens the article because it separates demonstrated deployment feasibility from deployment completion.

The same distinction applies to system latency more broadly. The chunked WatchConnectivity transfer improves robustness, but it also imposes a minimum delay that is independent of model complexity. The current system is therefore better described as near-real-time rather than interactive at the granularity of an individual footstep. That statement is important for scope control: the system already supports local inference and privacy-preserving operation, but it has not yet been characterized as a low-latency biofeedback platform.

### 6.3 Temporal Explainability and Reproducibility

Interpretability is treated here as a systems requirement rather than as a post-hoc add-on. Channel-level feature ranking answers which sensor matters. It does not answer when the signal matters within a stance phase. For health-related or biomechanical monitoring, that temporal question is essential because loading, mid-stance, and push-off represent mechanically distinct phases [11]–[13].

To address this, this work combines two complementary methods. Temporal SMILE masks contiguous 0.15 s windows within each input channel using a 0.10 s stride and measures the resulting change in model output [12]. A TimeSHAP-style eventwise method partitions each stance window into 20 equal bins, corresponding to 5% of normalized stance per bin, and estimates the marginal contribution of each segment [13]. The result is a channel-by-time explanation rather than a flat channel ranking. Figure 3(a) first shows the aggregate SHAP beeswarm view, which establishes that wrist channels dominate overall, with loading-to-push-off attribution ratios *×* of 4–6 for wrist acceleration channels and no waist channel in the top five.

**Fig. 3.**
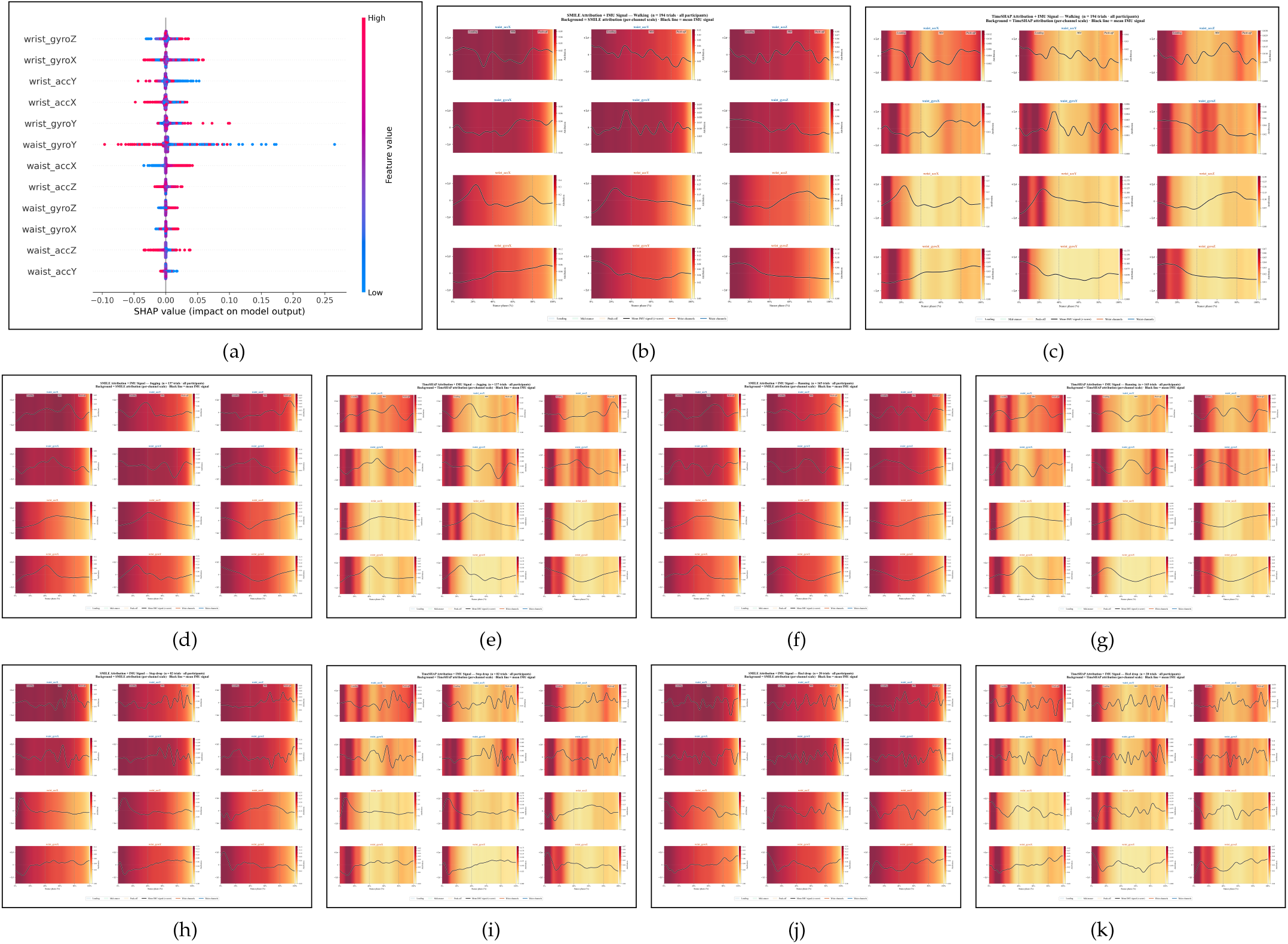
Temporal explainability and activity-specific attribution heatmaps. **(a)** Aggregate SHAP beeswarm across all 10 LOSO folds; wrist channels dominate the top ranks [11], [12], [13]. **(b)–(c)** Walking (*n*=137) Temporal SMILE and TimeSHAP: broad loading-phase wrist importance consistent with gradual heel-to-toe weight transfer; waist channels low-amplitude. **(d)–(e)** Jogging (*n*=136, *r*=0.867) Temporal SMILE and TimeSHAP: sharp loading-phase wrist peak; mild waist gyroscope mid-stance activation consistent with center-of-mass oscillation during cyclic gait. **(f)–(g)** Running (*n*=164, *r*=0.829) Temporal SMILE and TimeSHAP: wrist loading-phase dominance maintained; broader mid-stance waist gyroscope pattern consistent with two-peak GRF profile. **(h)–(i)** Step-down from 20 cm platform (*n*=82, *r*=0.442) Temporal SMILE and TimeSHAP: sharp wrist attribution spike at 0–15% stance; waist gyroscope mid-stance activation consistent with whole-body deceleration. **(j)–(k)** Heel drop (*n*=20, *r*=0.301) Temporal SMILE and TimeSHAP: narrow impulsive spike at 0–10% stance; biomechanically coherent pattern confirms correct event localisation despite lowest predictive accuracy. Ranking patterns remain stable across all 45 pairwise LOSO fold comparisons (median Spearman *ρ*=0.755, mean Kendall *τ* =0.589).

Both explanation methods converge on the same core result. Wrist acceleration during early stance emerges as the dominant reproducible signal. The attribution pattern is also physically coherent: heel strike propagates an impact up the kinematic chain, and the model appears to exploit that transmitted signature. The strongest part of the interpretability case is not the heatmap itself, but the stability analysis. Across all 45 pairwise LOSO fold comparisons, ranking patterns remain consistent, with median Spearman correlation of 0.755 and mean Kendall *τ* =0.589. `wrist_accX` appears in the top-three channels in every fold. That makes the explanation materially stronger than a one-off visualization from a single split and provides a mechanistic basis for the smartwatch-only fallback.

The comparison between Temporal SMILE and TimeSHAP is also informative in its own right. TimeSHAP concentrates 75.3% of attribution mass in the loading phase, whereas Temporal SMILE distributes attribution more progressively, with 50.4% of mass in loading. Rather than being contradictory, these differences reflect the distinct sensitivities of the two methods. TimeSHAP emphasizes coarse phase-level contribution, while Temporal SMILE resolves finer-grained within-phase structure. Their agreement on early-stance wrist dominance is therefore the most important result [12], [13].

### 6.4 Activity-Specific Behavior

The activity-level explanation matters because it links predictive performance back to movement mechanics. Figure 3(b)–(k) present paired Temporal SMILE and TimeSHAP heatmaps for all five activities. Together they show that the model does not fail randomly. It succeeds when the movement produces regular stance structure that is well represented in the filtered IMU stream, and it struggles when the target waveform is dominated by extremely brief impact transients.

Walking produces the broadest distribution of attribution across the stance phase. Informative signal is present across loading and mid-stance, reflecting gradual heel-to-toe weight transfer and the classic double-hump GRF profile. Jogging is the strongest activity in the dataset, achieving the highest mean correlation, and shows a sharper loading-phase wrist peak together with mild waist gyroscope involvement during mid-stance. Running preserves the cyclic regularity that benefits the model but shifts the explanatory pattern slightly, with broader mid-stance waist gyroscope activation consistent with the two-peak structure of running GRF.

Step-down is mechanically different from cyclic locomotion and this difference appears clearly in both performance and attribution. The explanatory spike is more onset-confined than in walking or running, and the waist gyroscope becomes more active through mid-stance, consistent with whole-body deceleration after landing from the platform. Heel drop is the clearest example of a biomechanically coherent but practically unsupported task. Both methods show a narrow impulsive attribution spike confined to the first 10% of stance, which matches the abrupt impact of dropping from standing height. The model is therefore attending to the correct physical region of the signal even though the event remains too brief for reliable waveform reconstruction under the present sensing and preprocessing pipeline.

### 6.5 Practical Utility for Monitoring

In many practical settings, end users do not need the full GRF waveform. They need summary metrics that can support comparison over time. The derived-metric analysis is therefore highly relevant to the wearable-computing value proposition. Impulse (*r*=0.983) and effective contact duration (*r*=0.982) agree very strongly with ground truth, making the system well-suited to longitudinal load tracking. Peak vGRF (*r*=0.741) and loading rate (*r*=0.588) require greater caution because they depend on narrow temporal windows where waveform errors are amplified.

The practical implication is direct. The present system is better suited to relative longitudinal monitoring than to exact peak-force assessment. Integrated measures tolerate local waveform error because deviations partly cancel over time. Peak-based and slope-based measures depend on narrow temporal regions and are more sensitive to small reconstruction errors or timing shifts. That distinction sharpens the article’s applied claim: the system is useful for repeated trend monitoring during cyclic locomotion, not as a drop-in replacement for every force-plate variable.

## 7 Limitations and Next Steps

The study has several limitations. The cohort is modest and limited to healthy adults, so the present results support subject-independent generalization within this dataset rather than broad clinical generalization. Stance segmentation in evaluation depends on force-plate timing, whereas a fully deployable wearable-only system will require IMU-based contact detection. The data were collected in a controlled laboratory environment, and generalization to outdoor terrain, footwear variation, and clinical populations remains untested. On-device timing and energy behavior also remain unmeasured.

These constraints define the current deployment scope. The evidence supports an edge-first wearable GRF estimation framework for cyclic locomotion under controlled sensing conditions. Phone-side latency and battery profiling on target hardware, IMU-only stance detection for end-to-end wearable operation, and field validation across surfaces, speeds, and patient populations remain important next steps.

## 8 Implications for Wearable Computing

The broader significance of the study extends beyond GRF estimation. The article illustrates a reusable design pattern for wearable AI: commodity body-worn sensing, local phone-side processing, compact temporal inference, privacy-preserving data locality, and reproducible time-resolved explanation. That pattern is relevant to many wearable applications in which connectivity is intermittent, data are sensitive, and users need outputs that can be understood and trusted.

This broader framing matters because the contribution is not only that the system predicts a biomechanical waveform. It is that the article connects device architecture, sensing, communication behavior, preprocessing, edge inference, explainability, and deployment boundaries in one coherent pipeline. That is what makes the work relevant to a wearable-computing audience rather than only a biomechanics audience.

## 9 Conclusion

This article presented an edge-first wearable system for estimating vertical ground reaction force from consumer smartwatches. Two Apple Watches stream inertial data to an iPhone, where preprocessing, storage, and inference are designed to execute locally without cloud dependence. The GRFNet-MultiScale model combines multi-scale dilated temporal convolutions with a global context branch to capture stance-scale dynamics in a compact architecture suitable for mobile deployment. Under strict LOSO evaluation, the dual-sensor system achieved a mean Pearson correlation of 0.798 with an RMSE of 257 N, while a wrist-only configuration retained 82.5% of dual-sensor correlation. These results establish that the signal required for useful vGRF estimation is present not only at the waist but also, to a meaningful degree, at the wrist.

The article also contributes an interpretability result that is more substantial than a single explanatory visualization. Temporal SMILE and TimeSHAP both identify early-stance wrist acceleration as the dominant signal, and that conclusion remains stable across validation folds. This matters because wearable systems intended for health-related use need more than prediction accuracy. They need a defensible account of what the model is using, when in the step it is using it, and whether that explanation persists when the train-test split changes. In that respect, the explainability analysis supports the broader trustworthiness of the system and provides a mechanistic rationale for the smartwatch-only fallback.

At the same time, the article draws a clear capability boundary. The present evidence supports cyclic locomotion as the strongest deployment region, with useful performance for walking, jogging, and running and more caution for step-down tasks. It does not support heel-drop-like impulsive events, fully characterized on-device execution, wearable-only stance detection, or broad field validation outside the laboratory. Those omissions should not be treated as minor caveats. They define the difference between an edge-ready research prototype and a clinically or commercially mature deployed system.

The most credible conclusion is therefore also the most useful one for the field. Consumer wearables can already support meaningful edge-first estimation of vertical GRF under controlled sensing conditions, provided that claims remain aligned with what has actually been measured. The next steps are clear: validate latency and energy use on target hardware, close the loop with IMU-only stance detection, and test generalization across surfaces, footwear, and clinical populations. If those steps are completed, the broader implication is significant. Force-plate-quality information will still belong to the laboratory, but repeated load monitoring between laboratory visits may increasingly be delivered by interpretable edge AI running on devices people already wear.

## Acknowledgments

This research was conducted at the University of Hull with ethical approval (FHS-24-25.036). We thank all study participants. The dataset is publicly available on Zenodo [15] under CC-BY-4.0.

---

**Parvin Ghaffarzadeh** is a PhD researcher in the Department of Computer Science at the University of Hull, Hull HU6 7RX, UK. Her research interests include edge AI for wearable biomechanics, deep learning for time-series data, and explainable AI for clinical applications. Contact her at.

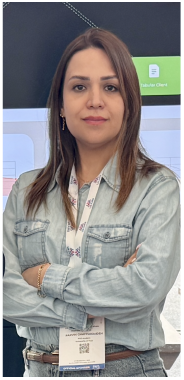

**Debarati Chakraborty** is a researcher in the Department of Computer Science at the University of Hull, Hull HU6 7RX, UK. Her research interests span machine learning, human motion analysis, and health informatics.

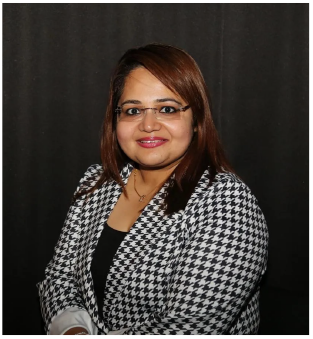

**Koorosh Aslansefat** (Senior Member IEEE),is an assistant professor of computer science at the University of Hull, HU6 7RX, Hull, UK. He is Co-Lead of the Dependable Intelligent Systems Centre and Deputy Lead of the Responsible AI Research Centre. In the past, he won prestigious research awards such as the IET Leslie H Paddle and Alan Turing Postgraduate Research. His research interests span artificial intelligence safety, Markov modelling, and real-time dependability analysis. Aslansefat received his PhD in computer science from the University of Hull. Contact him at

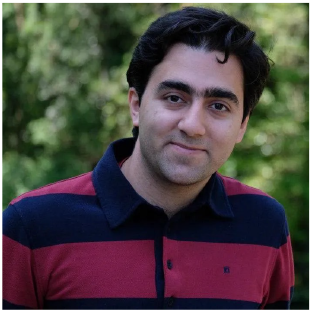

**Ali Dostan** is a lecturer in the Department of Engineering, Nottingham Trent University, Nottingham NG11 8NS, UK. His research interests include wearable sensing, clinical biomechanics, and rehabilitation engineering.

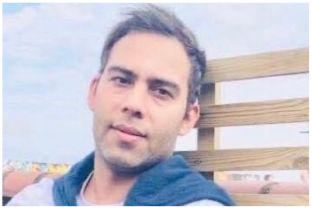

**Yiannis Papadopoulos** (Member, IEEE) is a professor in the Department of Computer Science at the University of Hull, Hull HU6 7RX, UK. His research interests span safety analysis, AI dependability, and autonomous systems.

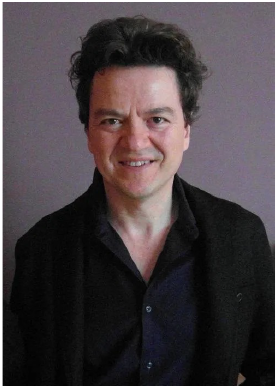

## References

[1] P. R. Cavanagh and M. A. Lafortune, “Ground reaction forces in distance running,” J. Biomech., vol. 13, no. 5, pp. 397–406, 1980.

[2] A. A. Zadpoor and A. A. Nikooyan, “The relationship between lower-extremity stress fractures and the ground reaction force,” Clin. Biomech., vol. 26, no. 1, pp. 23–28, 2011.

[3] W. B. Edwards et al., “Monitoring and managing load in sport,” Br. J. Sports Med., vol. 52, no. 23, pp. 1513–1514, 2018.

[4] D. A. Winter, Biomechanics and Motor Control of Human Movement, 4th ed. Hoboken, NJ: Wiley, 2009.

[5] E. Dorschky et al., “CNN-based estimation of sagittal plane walking and running biomechanics from measured and simulated inertial sensor data,” Front. Bioeng. Biotechnol., vol. 8, p. 604, 2020.

[6] W. R. Johnson et al., “Multidimensional ground reaction forces and moments from wearable sensor accelerations via deep learning,” IEEE Trans. Biomed. Eng., vol. 68, no. 1, pp. 289–297, 2021.

[7] M. Pogson et al., “A neural network method to predict task- and step-specific ground reaction force magnitudes from trunk accelerations during running,” Med. Eng. Phys., vol. 78, pp. 82–89, 2020.

[8] F. J. Wouda et al., “Estimation of vertical ground reaction forces and sagittal knee kinematics during running using three inertial sensors,” Front. Physiol., vol. 9, p. 218, 2018.

[9] R. S. Alcantara et al., “Predicting continuous ground reaction forces from accelerometers during uphill and downhill running,” PeerJ, vol. 10, e12752, 2022.

[10] M. Mundt et al., “Prediction of ground reaction force and joint moments based on optical motion capture data during gait,” Med. Eng. Phys., vol. 86, pp. 29–34, 2020.

[11] C. Rudin, “Stop explaining black box machine learning models for high stakes decisions and use interpretable models instead,” Nat. Mach. Intell., vol. 1, no. 5, pp. 206–215, 2019.

[12] A. A. Ismail et al., “Benchmarking deep learning interpretability in time series predictions,” in Proc. NeurIPS, vol. 33, pp. 6441–6452, 2020.

[13] J. Bento et al., “TimeSHAP: Explaining recurrent models through sequence perturbations,” in Proc. KDD, pp. 2565–2573, 2021.

[14] S. Bai, J. Z. Kolter, and V. Koltun, “An empirical evaluation of generic convolutional and recurrent networks for sequence modeling,” arXiv:1803.01271, 2018.

[15] P. Ghaffarzadeh, D. Chakraborty, K. Aslansefat, A. Dostan, and Y. Papadopoulos, “A multi-modal dataset for ground reaction force estimation using consumer wearable sensors,” Scientific Data, 2026. doi: 10.1038/s41597-026-07183-6. Dataset: doi:10.5281/zenodo.17376717.

